# Analyzing time-aggregated networks: the role of bootstrapping, permutation, and simulation

**DOI:** 10.1101/562231

**Authors:** T.R. Bonnell, C. Vilette

## Abstract

Networks are often used to describe adaptive social systems, where individual (node) behaviour generates network-level structures that influence subsequent individual-level behaviour. To address questions about the dynamics of network structure in these systems, there is a need to analyze networks through time. Various statistical methods exist for estimating the behaviour of networks in time, in terms of both time-ordered and time-aggregated networks. In this paper, we discuss three main analytical steps for the analysis of time-aggregated network data: 1) aggregation choices, 2) null-model comparisons, and 3) constructing, parameterizing, and making inferences from time series models. We then present a custom R package, netTS, which facilitates these steps. Observed grooming data from a group of vervet monkeys, a highly social primate species, is used as an example to highlight three potential analyses: 1) quantifying the stability of network-level social structures through time, 2) identifying keystone nodes driving/maintaining network structures, and 3) quantifying the interdependence between node behaviour through time. In particular, we highlight the role of bootstrapping, permutation, and simulation as critical components in the analysis of time-aggregated networks.

## 1 Introduction

Understanding the dependence between individuals in social groups has been enhanced by the use of network approaches. A network approach deconstructs a social group into nodes and edges, representing, respectively, individuals and the relationship between individuals (Newman 2010). This allows for the description of social dependence between two individuals (dyad scale), around the individual (ego scale), and at the entire group level (network scale). The network approach has become a successful and common method in many research fields, with the result that many theoretical and empirical predictions are tied to measurements of networks (Griffin & Nunn 2012; Chapman *et al.* 2016; Duboscq *et al.* 2016). In some cases these theoretical and empirical concerns are temporal in nature (Aplin *et al.* 2015; Formica, Wood & Brodie III 2017). For example, the magnitude of repeatability in social network position has direct implications for the selection of social phenotypes within a population (Aplin et al 2015). Similarly, in social inheritance models, the magnitude of a juvenile copying their mother’s social partners can have important consequences for the long-term stability of a population’s social structure (Ilany & Akçay 2016; Jarrett *et al.* 2018). Thus network measurements in time can have important implications.

Methodologically, the time-ordered and time-aggregated network constructions have been introduced for analyzing dynamic networks (Blonder *et al.* 2012). Time-ordered networks are networks that retain the timing of each interaction. These network constructions have been shown to be especially valuable when interested in questions about flow on a network (e.g., information, disease), as the timing of individual interactions can have important implications for the transmission between distant individuals (Blonder & Dornhaus 2011). Time-aggregated networks are networks constructed by aggregating data within a period of time, and can be useful for addressing questions regarding network topology. These networks lose the ability to directly query and use who contacted who and when, but can provide information about the structure of the network through time.

There are a variety of software packages that enable the analysis of networks in time. In particular, there are the timeordered (Blonder & Dornhaus 2011) and networkDynamic (Carter T. Butts *et al.* 2016) packages that handle time-ordered network construction, and can be used to extract time-aggregated networks. The purpose of the netTS package is to ease the analysis of time-aggregated networks by: 1) facilitating window sizes choices by comparing i) how the variation in the resulting time-series changes with window size, as well as ii) how the uncertainty in a network measures over time changes with window-size using bootstrapping, 2) contrasting the observed time series to null models using network permutations, and finally, 3) simulating temporal relational data to test, refine, and interpret statistical models used to analyze time-aggregated networks.

In this paper we first give an overview of analyzing time-aggregated networks using the netTS package, providing details of its use, followed by examples that highlight how it is possible to address questions concerning the temporal dimensions of social structure.

## 2 Methods

To introduce the netTS package, we first present the moving window approach for aggregating relational data. We then address options for window size choices, estimating network measures through time, as well as for the analysis of these measures, once extracted. The full package code can be found on github (github.com/tbonne/netTS), along with tutorials and the code used in the analysis presented in this paper.

After this introduction to the package, we present examples showcasing the abilities of the method to take advantage of both network and time-series analysis. We first show how one can assess network stability through time, detect and quantify keystone nodes using generalized additive mixed models, and finally, how it is possible to estimate the dependence in social behaviour through time between nodes using multivariate autoregression (MAR). We also demonstrate how simulated data can help guide inferences and parameterization of statistical models by simulating data for the keystone and social dependence analyses. These examples are meant to illustrate how combining social network and time series approaches can facilitate the analysis of social questions that are inherently temporal in nature.

### 2.1. The Moving window approach to aggregation

Generally, when constructing social networks using time-aggregated networks to interrogate relational data, some amount of aggregation is required, e.g., grouping all data into years to create yearly networks (Blonder *et al.* 2012). This package aims to help with this process using a moving window approach.

The moving window approach allows a user to define its size (e.g., windowsize = 1 month) and the amount to shift the window (e.g., windowshift = 1 day). First, this moving window subsets the relational data within a window and creates a network. It then shifts in time and repeats the process. By altering its size and shift, it is possible to generate a time series of networks (Fig. 1), and can be thought of as generating a multilayered network in which each network layer encodes the same type of interactions at different time points (Finn *et al.* 2019).

**Figure 1:**
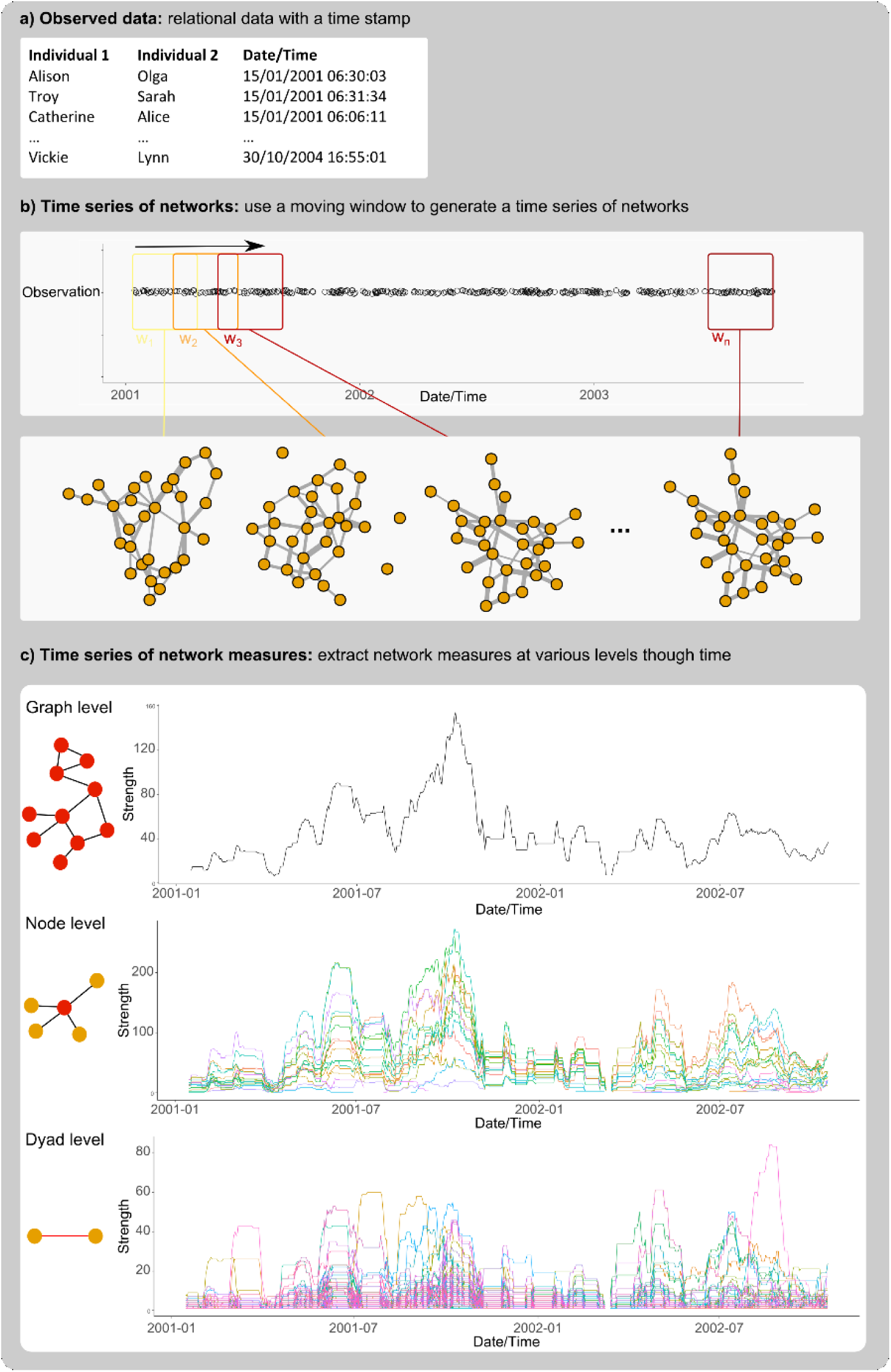
Overview of using a moving window approach to extract network measures over time: a) the relational input data, b) generate the time series of networks using a moving window approach, and c) extract network measures at the dyadic, node, and network scales.

### 2.2. Window size choices

The ability to alter the window size and shift introduces the possibility of multiple scales being chosen. With the possibility of longer window sizes, more data is aggregated together within each network. The lower limit to window size choices can, to some extent, be specified by the fact that as the size gets too small, the network measures become progressively noisier (see the bootstrap method discussed below). The upper limit however, apart from the maximum time scale of the dataset, has no *a prior* limit and will increasingly capture longer-term trends. In some cases, depending on the temporal dynamics of the systems of interest, there are potentially “natural” scales (Caceres, Berger-Wolf & Grossman 2011). In general however, apart from attempting to identify one optimal window size, it is likely the case that the way a pattern of interest changes depending on the time scale chosen will itself be of great interest.

The amount to shift a window through time will affect the scale at which change occurs, with smaller window shifts capturing shorter time-scale changes. Similarly, smaller window shift choices will also increase the amount of autocorrelation in the time series, as subsequent window aggregations will share much the same relational data. Depending on the method being employed to analyze these data, this can present a challenge, as some statistical methods are better at modeling or accounting for temporal autocorrelation.

### 2.3. Network measurement through time

#### 2.3.1. Extracting measures

Once a window size and shift has been selected, and a time series of networks generated, it is possible to use network metrics at the scale of the network, node, or dyad (Newman 2010). A few common metrics are built in to the netTS package, but in general the network measure required is a user-specified function. This function takes a network as input and returns a value, or vector of values, in the case of node or dyadic measures (e.g., Fig. 2). By using user-generated functions, the package can take advantage of the wide range of network measures available, without constraining users to a pre-specified list of options.

**Figure 2:**
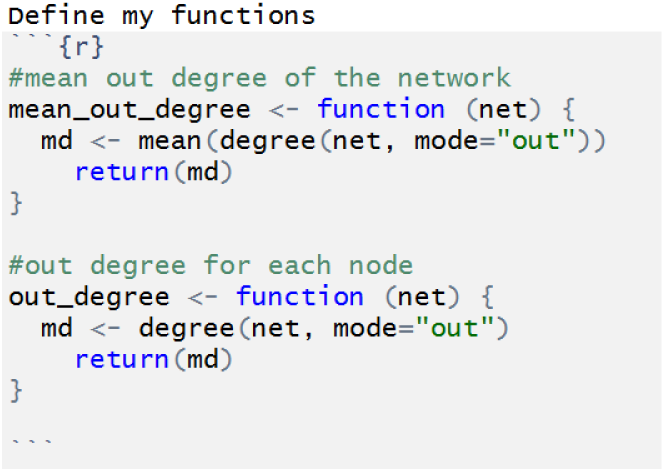
Example of user-defined functions. The first function takes a network as input and returns one value for the whole network; the second function takes a network as input and returns a value for each node.

#### 2.3.2. Controlling for sampling effort/time

Given the time series nature of the data, as well as the potential for variation in data collection methods through time, it is important to consider how changes in sampling effort might impact a potential measure. In some cases, the ability to use scaled or transformed measures such as the simple-ratio index (SRI) can facilitate comparisons between networks in time (Farine & Whitehead 2015). Another option, that keeps the measure on the observed scale, would be to directly control for sampling effort over time. This converts the observed social behaviour to a rate (e.g., interactions/hours of observation). To facilitate this conversion in netTS, it is possible to include a sampling effort function. It takes as input an event’s dataframe and returns a single value of effort. The netTS package provides two functions. The first approach sums the total time (effort.time), based on the first and last sample time of each day within a window. This method assumes equal sampling effort throughout the day and could be useful for data collected ad libitum throughout a sampling day. The second approach uses a unique ID number for each scan event (effort.scan). It assumes that events are captured within set scanning periods, with each period given a unique ID, and would be suitable for sampling regimes where periodic scans are used to collect data. As sampling effort will vary by dataset and collection method, it is also possible to construct user defined effort functions.

#### 2.3.3. Network measure accuracy

To assess the accuracy of a network measure, we use a bootstrap approach on the event data used to create the network. Applying this method, it is possible to take multiple bootstrap samples of the event data within a time aggregated window, create a network with the bootstrapped sample, calculate a network measure, and then estimate the correlation between measures in the bootstrapped networks and the observed network. Higher correlation estimates indicate that the network measure is robust to bootstrapped sampling, suggesting that sampling is adequate to provide a good measurement. This test can be advantageous for detecting the lower threshold for the window size choices.

### 2.4. Analyzing measures through time

#### 2.4.1. Network null model comparisons

Given the ability to compare how a network changes in time, it can also be useful to contrast how this changing network compares with a null model using network permutations. The exact specification of the null model, i.e., how it is constructed, can aid in understanding the structure of the observed network. For example, it is possible to construct a time series of centrality measures within a grooming network and look for trends over time. However, if we want to compare that centrality measure to what might be expected due to grooming interactions driven by chance alone, we would want to use a null model. Here, the choices of the null model can help refine how the observed pattern is different (Farine 2017). We could decide to take all grooming events and randomly distribute them between nodes to generate a null model. Similarly, we could maintain that some individuals are more present in grooming events than others by permuting individuals between grooming events. We can then compare the observed network to those null models to make inferences about how it differs or not. By performing permutations for each time-aggregated network, it allows for estimations of how the network diverges from the null model through time (e.g., is it consistently different, or are there only certain times/seasons where there is difference?). In the netTS package, we provide some predefined permutation methods, but also allow for user-specified permutation functions that will take an event data frame as input and return a range of network measurement values.

#### 2.4.2 Use of simulated data

Simulated data can often be crucial in suitably selecting a modeling framework, parameterizing the chosen model, and correctly interpreting the model results. In the netTS package, we offer a simulation function to generate an event data frame, the output being a set of interactions between a set of nodes with an associated time stamp. The simulation models change in the probability of an individual performing an interaction with another individual, and constrains with whom the individual interacts using a specified network. The probability model is of the form:

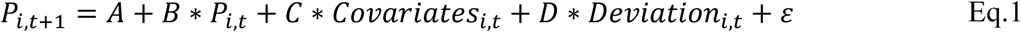

Where *P*_*i,t*_ is the probability of individual *i* performing an interaction at time *t*, A is a vector of mean probabilities for each individual, B is a correlation matrix describing the dependence between individual probabilities, C is a matrix of effects of covariates on the probability of each individual, D is a correlation matrix describing the dependence between individual deviations from their mean values (i.e., A), and ε is a scale parameter which controls the amount of noise. At each time step node, probabilities are updated and a stochastic sample is taken from the population. If a node is selected to perform an interaction during a sampling event, it randomly selects one out of its neighbour in a pre-specified social network.

At its base, the simulation function can be used by simply choosing the number of nodes, the number of scans to take, and the number of scans per day. In this case, as A, B, C, D, ε, and the underlying network are not specified, a set of defaults values are used to produce an event data frame based on a random grooming network and fixed individual behaviour within no inter-individual interactions. From this simple simulation case, it is possible to add a known network, interdependence between node behaviour, as well as covariates effects. The complexity of the simulation will largely be driven by the goal of the analysis: e.g., can the statistical method correctly identify no pattern (random simulation), or identify a known pattern (structured simulation)?

## 3. Using netTS: an example of a primate social network

### 3.1 Input data

We use grooming data from a fully habituated group of vervet monkey in the Eastern Cape of South Africa (Josephs *et al.* 2016). These gregarious primates occupy a semi-arid environment with large seasonal fluctuations of both temperature and rainfall, and similarly show seasonal breeding patterns (Lubbe *et al.* 2014; McFarland *et al.* 2014; McFarland *et al.* 2015). The grooming data consists of observations of grooming events, e.g., Laur grooms Malc @2015-07-01 12:32:19.

**Table 1:** example data used as input for the netTS package.

| **from** | **to** | **date** |
| --- | --- | --- |
| <chr> | <chr> | <S3: POSIXct> |
| Laur | Malc | 2015-07-01 12:32:19 |
| Laur | Malc | 2015-07-01 12:32:19 |
| Malc | Laur | 2015-07-01 12:33:01 |
| Malc | Laur | 2015-07-01 12:33:01 |
| Ubun | Wall | 2015-07-01 16:08:26 |

### 3.2. Correct for changing sampling effort

Given that sampling effort can vary between time periods, it is important to control for it when comparing certain network measures over time, with some measures being more sensitive to sampling effort than others: e.g., node strength vs. node degree. Here, we demonstrate how controlling for time spent sampling in the field alters average strength of a network over time (Fig. 3).

**Figure 3:**
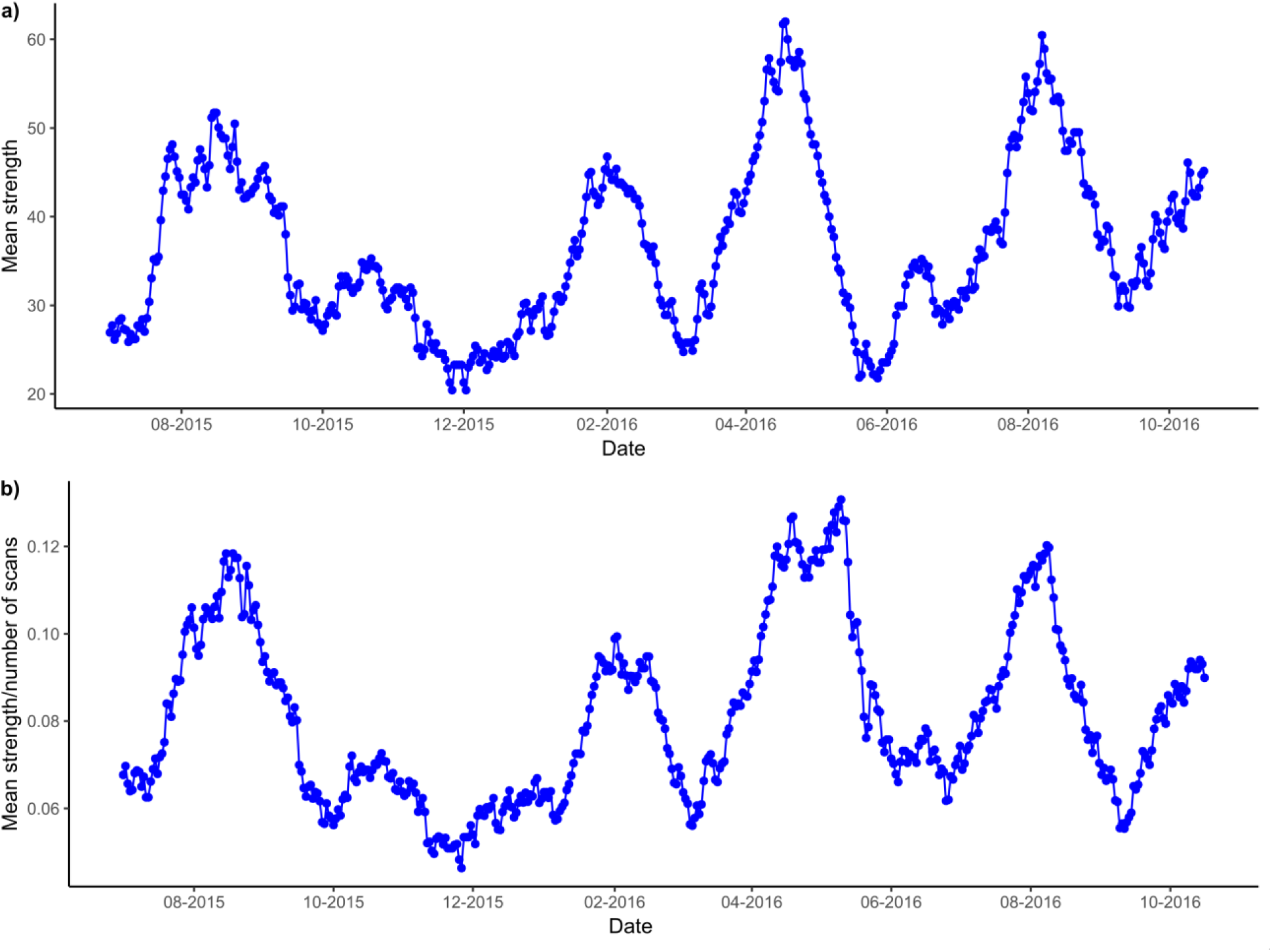
Mean network strength over time for a group of vervet monkeys: a) without correcting for sampling time in the field, and b) after correcting for sampling effort.

### 3.3 Assess window size choice

We first vary the window size from 10 days to 200 days to see how the variance in edge density changes. The idea here is that at small window sizes, the density will be consistently low, reducing variation. Similarly, at high window sizes, the density will be saturated, resulting again in low variation across time. We therefore look for window sizes that maximize the variation in edge density (Fig. 4). In this case variation in edge density showed a peak around 33 days (Fig. 4a).

**Figure 4:**
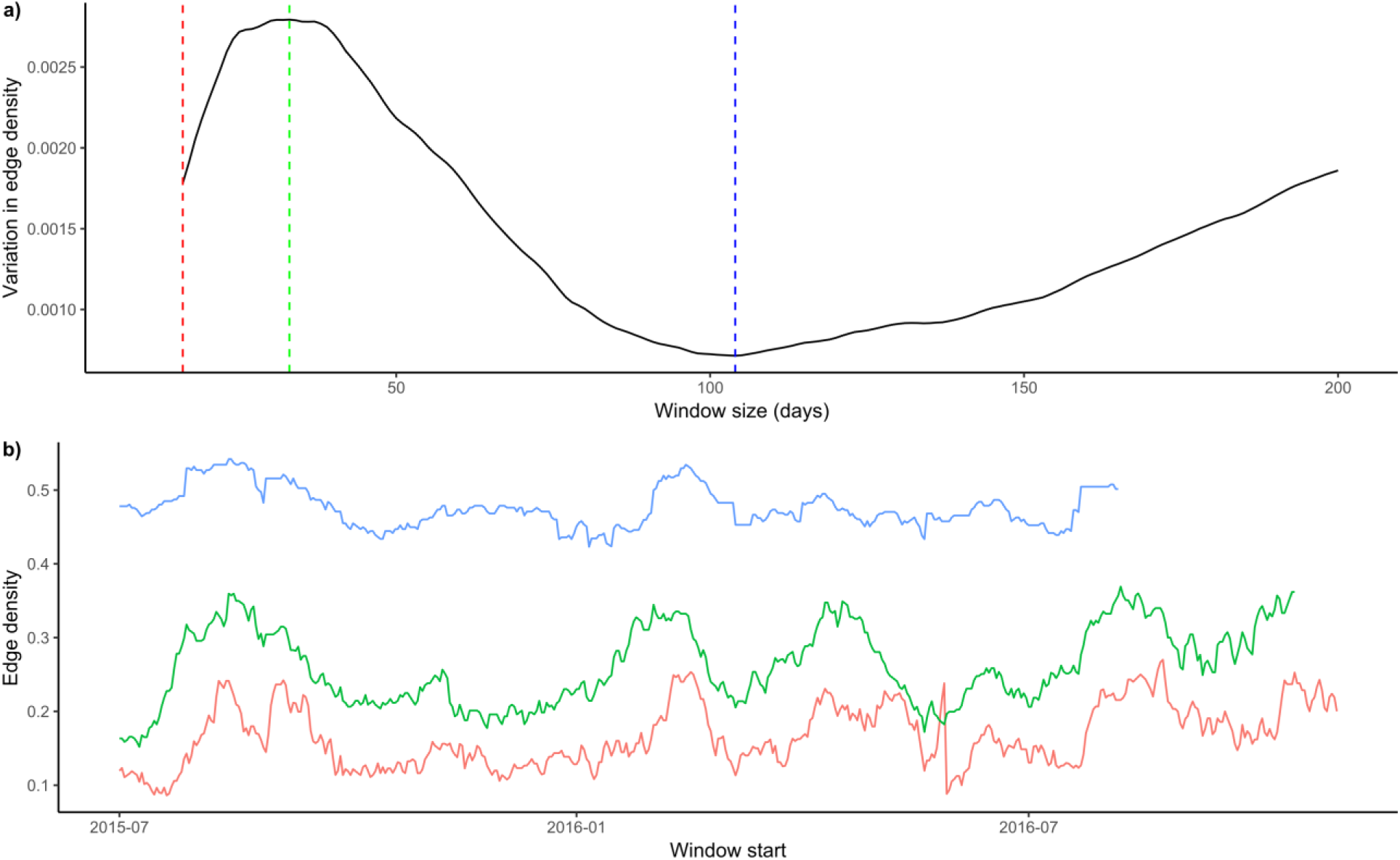
The relationship between window size choices and the density of edges in the network: a) how the variation in edge density changes with window size choice, and b) plot of edge density over time for widow size choices of 16 days (red), 33 days (green), and 104 days (blue). The colours of the vertical lines in a) correspond to colours of lines in b).

To highlight how window size choices impact subsequent network measures we plot mean node out-strength over time for a range of window sizes. We use the bootstrap test in order to identify the lower end of possible window size choice (Fig. 5). The results suggest that, given the temporal resolution of the vervet data network, measurement accuracy is reduced in window sizes below 30 days. Notably, the consistently high similarity between the bootstrapped networks and observed networks, using a 30-day window, suggests that the window size identified by the maximum variation in edge density (i.e., 33 days) results in robust networks.

**Figure 5:**
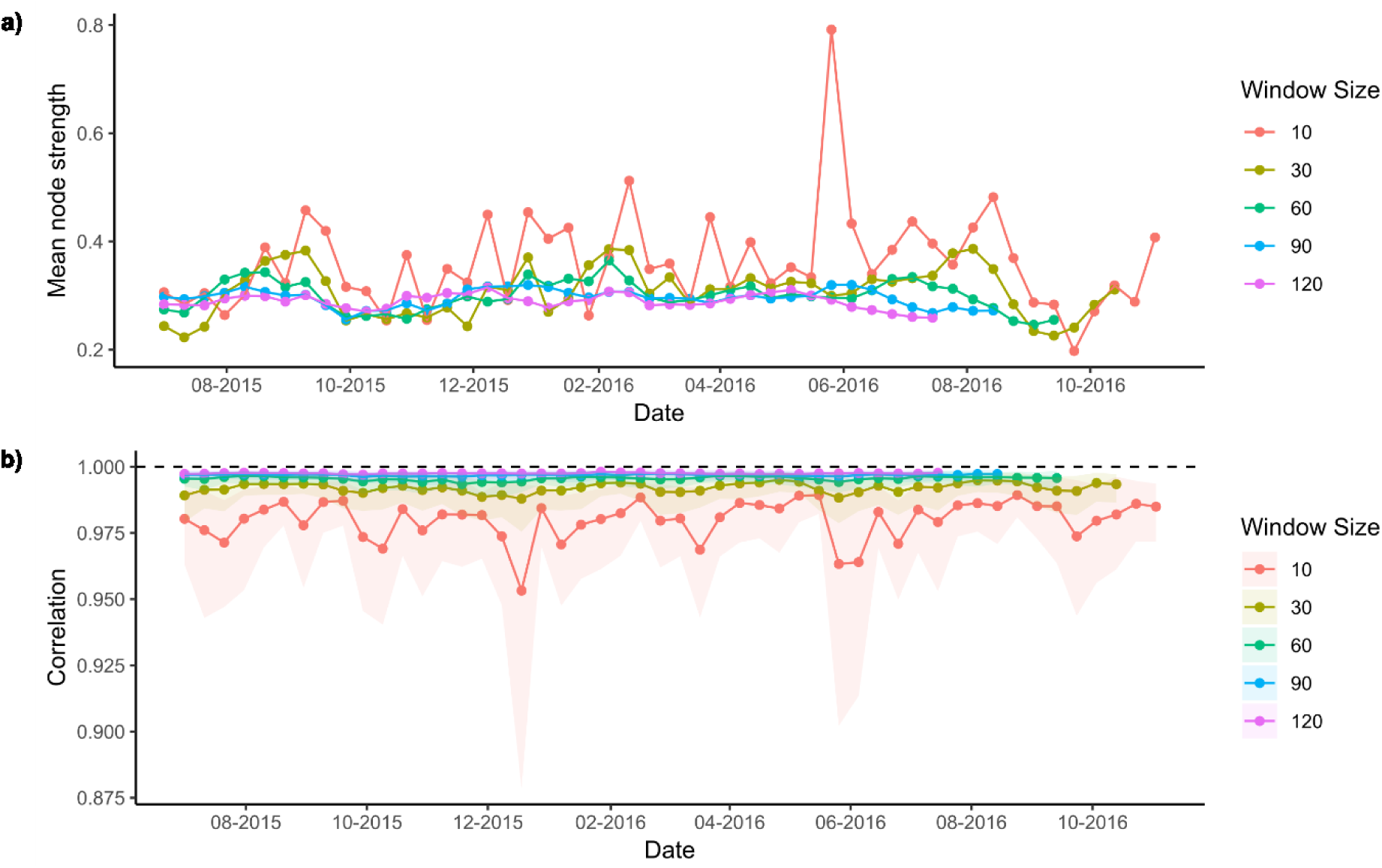
Results using different window size values, i.e., different aggregation choices: a) mean node strength under alternative window sizes, and b) correlation between node strength of the observed network and that of bootstrapped replicates. Higher mean correlation estimates and lower variability around these estimates suggest more robust network measurements.

### 3.4. Assessing network structure through time

To assess how social structure deviates from random through time, we use a permutation approach. Here, for each network in the time series, the observed measure is compared to a randomly permuted network. The package contains options for user-specified permutation functions giving users the ability to set the method of permutation up in order to address a particular question (Farine 2017). In this example, we randomly swapped individuals in the event data frame, mimicking random grooming while keeping the frequency with which individuals are seen involved in grooming the same. We then tested for differences between the pattern of “random” grooming and our observed grooming patterns through time. We used this approach to assess the consistency of mean out-degree (the number of partners groomed) and mean eigenvector centrality of the network (magnitude to which the grooming interactions concentrate on a few well connected individuals) (Fig. 6).

**Figure 6:**
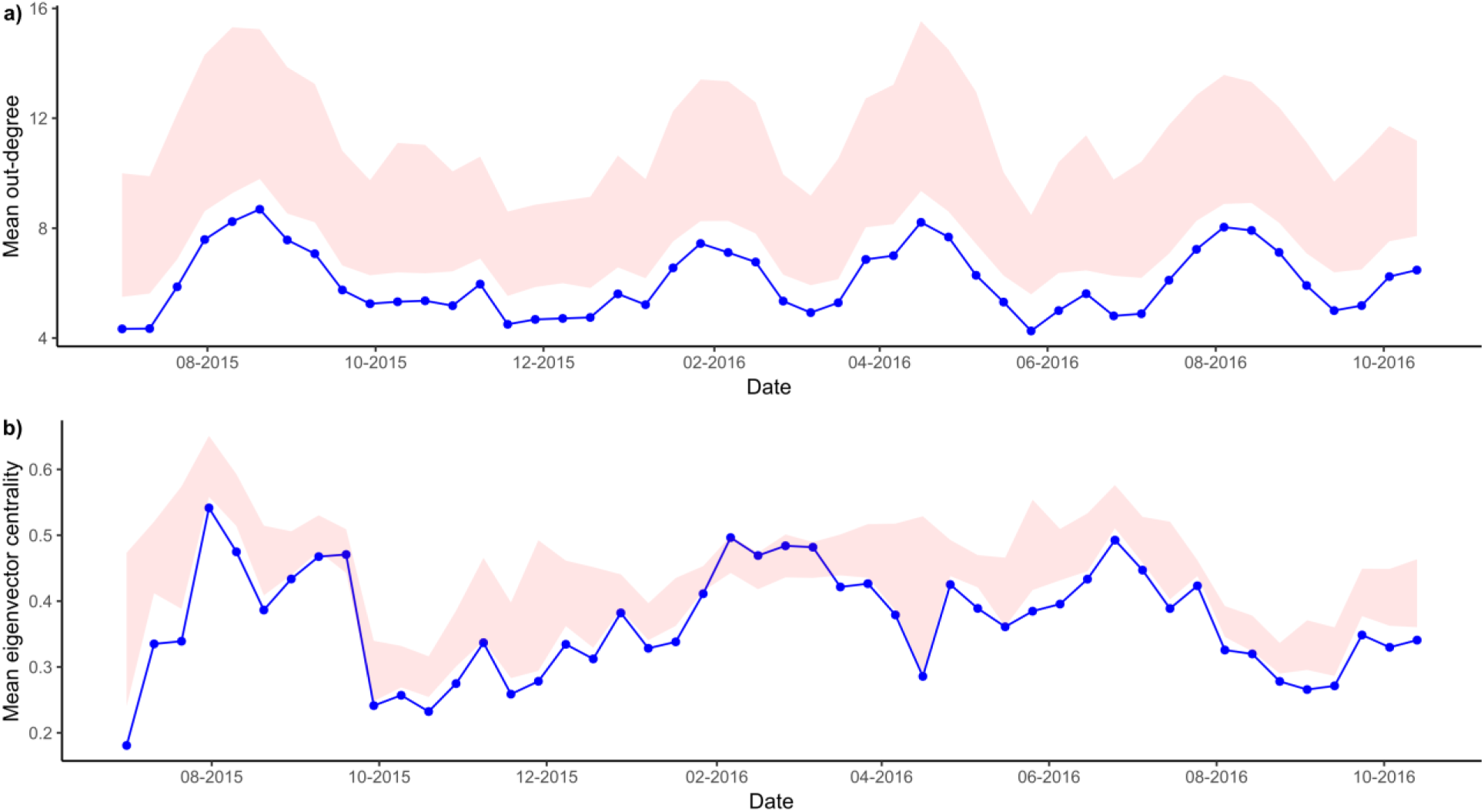
Permutation across time: a) mean out-degree of grooming, and b) mean eigenvector centrality. The observed values are presented as blue points, and the 95% quantiles generated through permutations are presented as a pink ribbon.

We can see from figure 6 that, within the group, out-degree is consistently lower than expected with random grooming interactions, i.e., that individuals are more selective with whom they groom compared to random. Whereas, in the case of group centrality, there is less differentiation between random and observed networks, with only occasional times when mean eigenvector centrality is not lower than expected by chance.

We can similarly evaluate the stability of both a particular network structure (e.g., mean out-degree, eigenvector centrality) as well as the network as a whole. To measure how much a network as a whole changes over time, we make use of cosine similarity (Newman 2010), a built-in function in the netTS package. We calculated both the magnitude to which the network differs each window shift, as well as the magnitude that the network has changed since the first observed network (Fig. 7).

**Figure 7:**
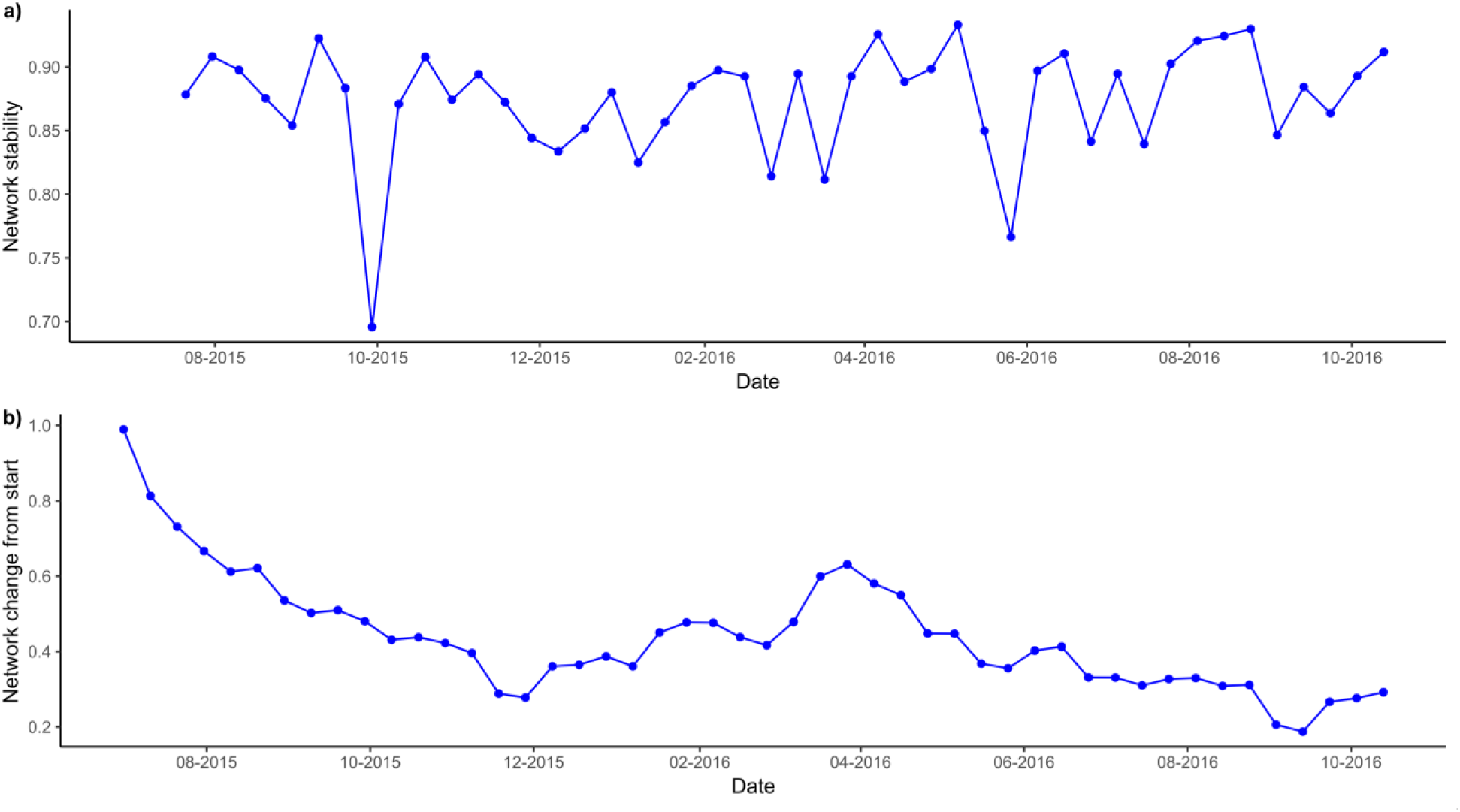
Change in network structure over time, measured as a) the change at each window shift (10 days), and b) the overall change in network structure compared to the initial network.

We found that in the case of grooming structure within the vervet monkey group, we see that the network remains similar between 10-day shifts, except for a noticeable spike around October 2015 (Fig. 7a). In the overall network structure, we also found a steady decline from the beginning of 2015, with a slight increase in similarity in early-2016 (Fig. 7b).

### 3.5. Identifying keystone individuals

To identify keystone individuals (i.e., nodes) we look to see how individual changes in out-grooming behaviour influenced the centrality of the group as a whole. Here, we are interested in answering the question: do some individuals influence the social structure of the group more than other individuals when they groom? With the netTS package, we extracted individual out-grooming strength, and eigenvector centrality of the network over time. We then used a generalized additive mixed model to estimate how changes in out-grooming influence the eigenvector centrality of the network. We allowed this effect to vary by individual by using a random slope for the effect of out-grooming. If this random slope turns out to be negligible, it would suggest that out-grooming behaviour for all individuals has the same effect on group centrality structure. We also control for seasonal effects via a circular basis spline on day-of-year, and model dependence in the residuals using an AR1 process. We fit the model with the brms package following a Bayesian approach (Bürkner 2017).

The model suggests that there are some differences between individuals in the effect of their grooming on centrality of the group (standard deviation in the effect of grooming: sd(grooming) = 0.49, 95%CI: 0.31,0.72). Running the model with and without a random slope (∆WAIC = −27.26, se = 24.70) suggests that there is evidence for individuals that are associated with increases/decreases in centrality when their out-grooming increases, and points to potential keystone individuals (Fig. 8).

**Figure 8:**
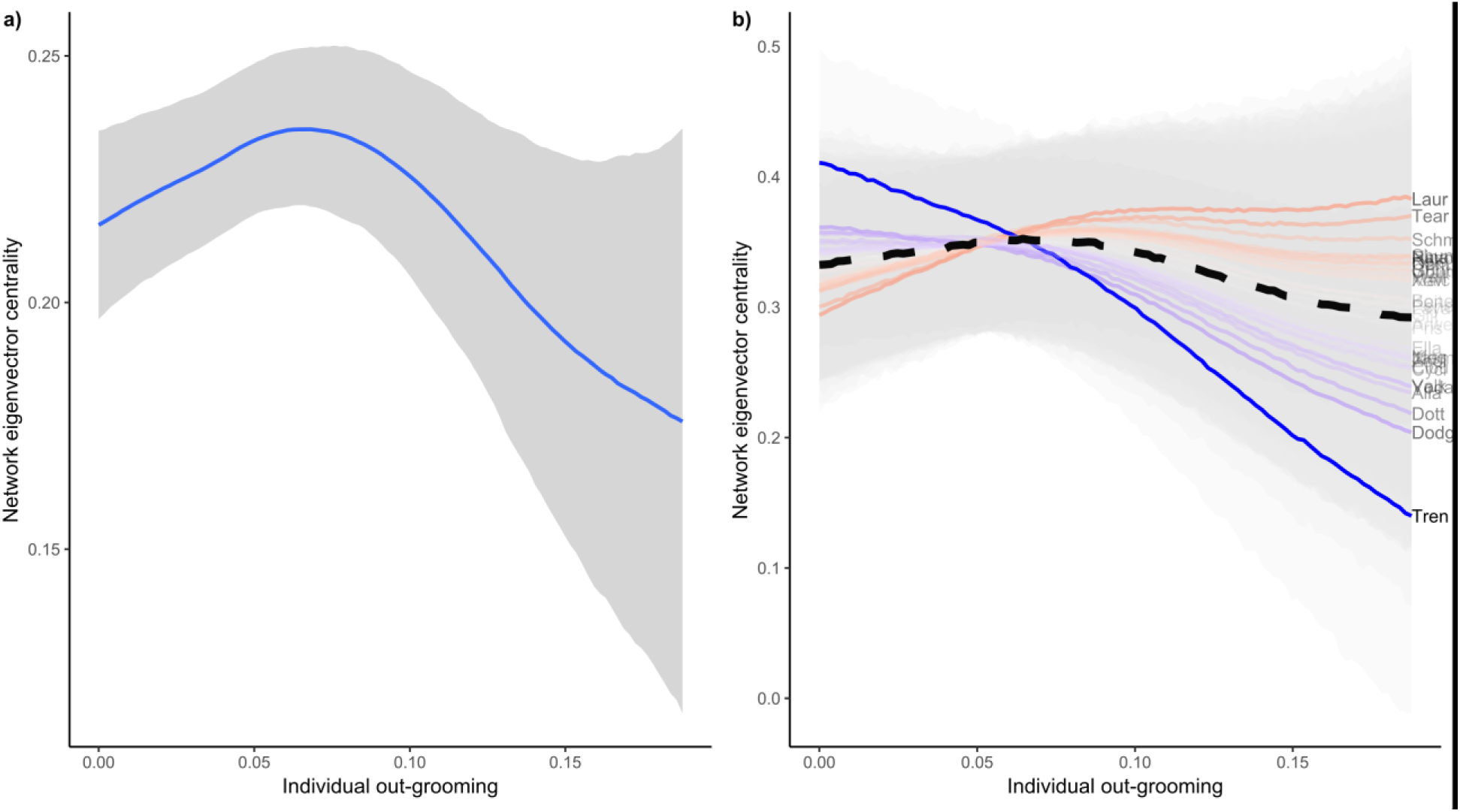
Model estimating how changes in out-strength by nodes influence the centrality of the network as a whole: a) overall mean effect of out-grooming on network eigenvector centrality, and b) individual level effect of out-grooming on network eigenvector centrality. Individual lines represent the effects of particular nodes (individuals) and are coloured based on their deviation away from the mean effect: i.e., blue is lower than the mean, and red is above the mean effect. Shading indicates the 95% credible intervals for each line. Each line is also given labels based on the name of the individual to aid in identifying those individuals having either a more positive or negative effect on eigenvector centrality.

We then use simulated data, using a known underlying grooming networks, to aid in making inferences from the results. Running the model on simulated data with a fully connected grooming network suggests that when individual grooming is not constrained, i.e., all individuals have equal probability of receiving grooming, the model found no difference in individual influence on group eigenvector centrality (sd(grooming) = 0.000, 95CI: 0.000,0.000). However, in the case where individuals show constraint in their grooming behaviour, and where some individuals are more central in the group than others, the model found differences in the effect of node out-strength on centrality of the group (sd(grooming) = 0.001, 95CI: 0.0005,0.010). In particular, those more central to the group tend to reduce centrality when they increase grooming, whereas more peripheral individuals tend to increase centrality (Fig. 9).

**Figure 9:**
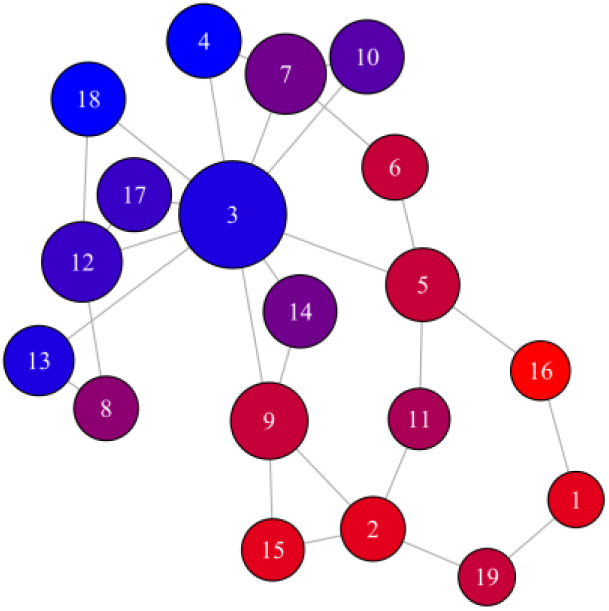
Effect of individual grooming behaviour on the centrality of the group, estimated from simulated data. Colour of nodes are set based on their deviation from the mean effect, with red nodes increasing centrality of the group with increased grooming and blue nodes decreasing centrality with increased grooming.

### 3.6. Quantifying social dependence

The ability to extract network measures as time series affords many analytical options. As an example, a multivariate autoregressive model (MAR, also known as vector autoregression) was used to estimate the dependency between individual measures of grooming behaviour in our vervet group (Ives *et al.* 2003). This allowed us to test whether individual changes in out-grooming behaviour are correlated among individuals, i.e., do some individuals show coordinated out-grooming behaviour.

We used simulated data to help construct and interpret the MAR model. We simulated two scenarios: 1) where individuals do not adjust their out-strength in response to others, and 2) where only two individuals adjust in response to changes from the other (correlation = 0.3). These two simulations allow for a test of the MAR in terms of how well it can accurately find no pattern when none exists, and how well it can find one when it does exist. We simulate data with the sim.events.data() function in the netTS package, and match the number of individuals (19) and the sample size of the observed data. For the first scenario, we fix each individual’s grooming behaviour between 0.1 and 0.9 using a uniform random distribution. For the correlation matrix D, we fill diagonal cells with 0.4, and off diagonal values are set to 0. This correlation matrix represents the case where there are no inter-individual correlations (off-diagonal), and medium within individual autocorrelation (diagonal) in deviations around the fixed mean. For the second simulation, we again use fixed means randomly selected between 0.1 and 0.9, but choose a correlation matrix D with the diagonals set to 0.7 and the off diagonals between one dyad set to 0.3. Once the simulated event datasets were constructed, we extracted out-grooming strength of individual nodes with the nodeTS function.

The results of the MAR run on simulated data without a pattern showed estimated inter-individual correlations ranging between (min = −0.28, max = 0.28), and did not find a correlation where the estimated 95%CI did not contain 0. The MAR model run with the dataset generated with a known pattern, estimated individual correlations between (min = −0.23, max = 0.50), and estimated the correlation of the known pair to be 0.50 (95%CI = 0.16, 0.81). The simulation tests suggested that given our data the MAR is likely to detect correlated out-grooming, however the results, given our statistical model and the data collected, are likely to be noisy. To help parameterize the model to make better inferences, the simulated data with no pattern was used to help choose priors, reducing the chance of false positives. Similarly, the simulation with a known pattern found that the best results were achieved when non-overlapping windows were extracted using nodeTS.

The results from the MAR model run on the observed data suggested only a weak interdependence between individual out-grooming behaviours, with positive correlation estimates of 0.31 between Laur and Tear and, 0.34 between Alla and Zool (Fig. 10). Interestingly, in the case of the Laur-Tear dyad, individually, they are also estimated to have a higher than average positive effect on the centrality of the grooming social network of the group. This suggests that when one increases/decreases grooming, so, too does the other, and that both lead to a disproportionate change in centrality of the grooming network. For all other dyads, the sign of the correlation was not certain using the range of the 95% credible interval.

**Figure 10:**
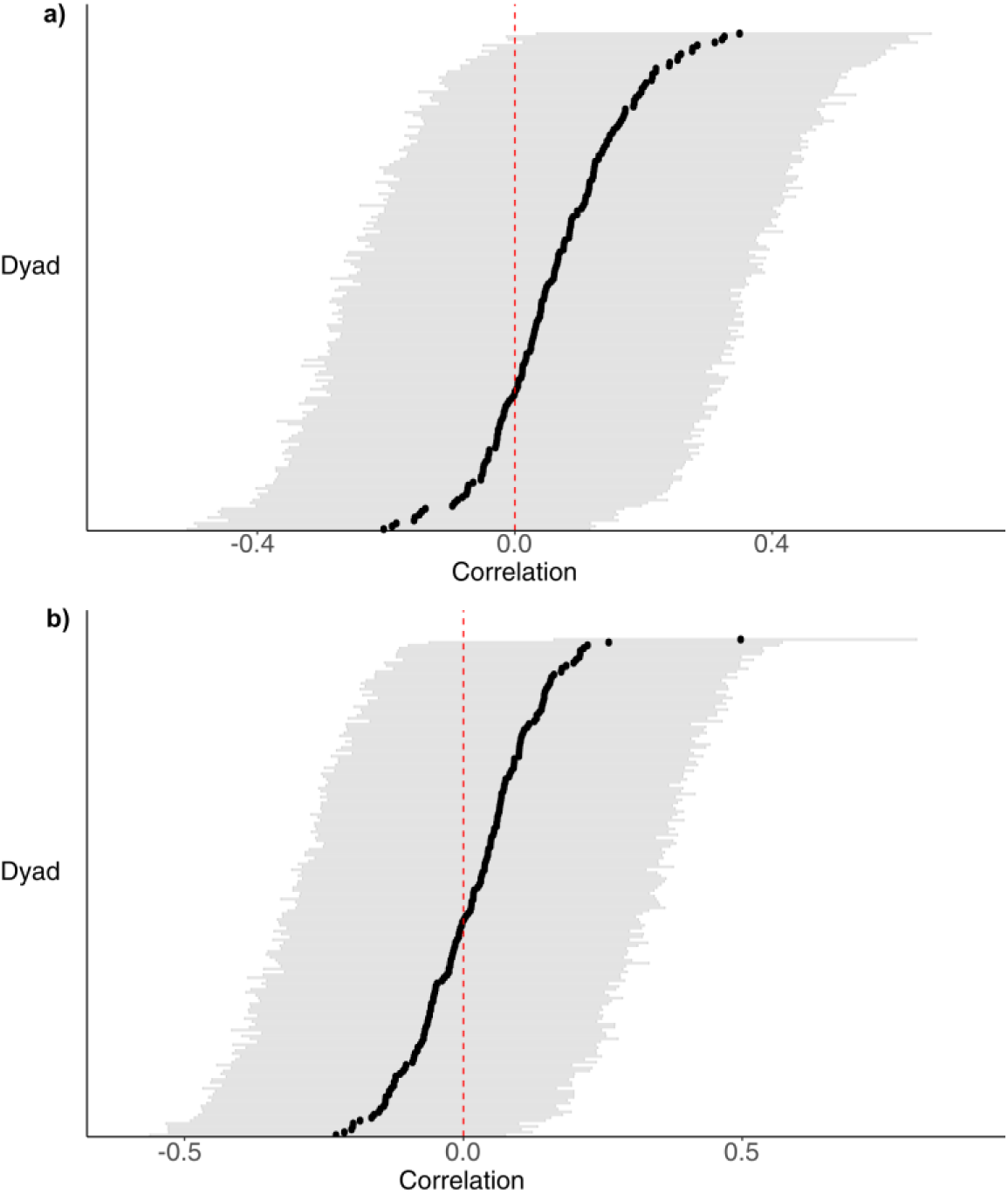
MAR estimate of interdependence in out-strength of grooming behaviour between all dyads: a) the cross-correlation estimates for the observed out-grooming behaviour, and b) the cross-correlation estimates for the simulated data with only one dyad having inter-dependent grooming behaviour. Black points represent mean correlation estimates, and grey lines indicate the 95% credible interval. The red dashed line highlights zero correlation between dyads.

## Conclusions

The construction of time-aggregated networks requires careful consideration of scale. Similarly, the choice, parameterization, and interpretation of statistical models employed to analyze the resulting time-series of networks require careful inspection. We suggest that the use of bootstrap, permutation, and simulation can facilitate decision-making regarding these choices, and have introduced the netTS package for this purpose.

## Appendix

List of vignettes available (github.com/tbonne/netTS):

1. Introduction
2. Choosing a window size
3. Controlling for sampling effort
4. Controlling for nodes entering and leaving the network
5. Network similarity
6. Using network permutations
7. Using network simulations

## Acknowledgements

We thank Mark and Sarah Tompkins for the permission to work at Samara, Kitty and Richard Viljoen for their continued logistic support, and Louise Barrett and Peter Henzi for insightful discussions and computational equipment. We are also very grateful to the many research assistants who contributed to the database, without which only simulated data would be available.

## Author’s Contributions

TRB and CV conceived the ideas and designed methodology; CV collected the data; TRB analysed the data; TRB led the writing of the manuscript. All authors contributed critically to the drafts and gave final approval for publication.

## Data Accessibility

All data used in the paper has been uploaded to github and is available from the package netTS (https://github.com/tbonne/netTS). Similarly, the code for all analyses are available at: https://github.com/tbonne/netTS/tree/master/inst/manuscript.

## References

Aplin, L.M., Firth, J.A., Farine, D.R., Voelkl, B., Crates, R.A., Culina, A., Garroway, C.J., Hinde, C.A., Kidd, L.R., Psorakis, I., Milligan, N.D., Radersma, R., Verhelst, B.L. & Sheldon, B.C. (2015) Consistent individual differences in the social phenotypes of wild great tits, Parus major. Animal Behaviour, 108, 117–127.

Blonder, B. & Dornhaus, A. (2011) Time-ordered networks reveal limitations to information flow in ant colonies. Plos One, 6, e20298.

Blonder, B., Wey, T.W., Dornhaus, A., James, R. & Sih, A. (2012) Temporal dynamics and network analysis. Methods in Ecology and Evolution, 3, 958–972.

Bürkner, P.-C. (2017) brms: An R package for Bayesian multilevel models using Stan. Journal of Statistical Software, 80, 1–28.

Caceres, R.S., Berger-Wolf, T. & Grossman, R. (2011) Temporal scale of processes in dynamic networks. Data Mining Workshops (ICDMW), 2011 IEEE 11th International Conference on, pp. 925–932. IEEE.

Carter T. Butts, Ayn Leslie-Cook, Pavel N. Krivitsky & Bender-deMoll, S. (2016) networkDynamic: Dynamic Extensions for Network Objects. R package version 0.9.0. https://CRAN.R-project.org/package=networkDynamic.

Chapman, C.A., Friant, S., Godfrey, K., Liu, C., Sakar, D., Schoof, V.A., Sengupta, R., Twinomugisha, D., Valenta, K. & Goldberg, T.L. (2016) Social behaviours and networks of vervet monkeys are influenced by gastrointestinal parasites. Plos One, 11, e0161113.

Duboscq, J., Romano, V., Sueur, C. & MacIntosh, A.J. (2016) Network centrality and seasonality interact to predict lice load in a social primate. Scientific Reports, 6, 22095.

Farine, D.R. (2017) A guide to null models for animal social network analysis. Methods in Ecology and Evolution, 8, 1309–1320.

Farine, D.R. & Whitehead, H. (2015) Constructing, conducting and interpreting animal social network analysis. Journal of animal ecology, 84, 1144–1163.

Finn, K.R., Silk, M.J., Porter, M.A. & Pinter-Wollman, N. (2019) The use of multilayer network analysis in animal behaviour. Animal Behaviour, 149, 7–22.

Formica, V.A., Wood, C. & Brodie III, E. (2017) Consistency Of Animal Social Networks After Disturbance. Behavioral Ecology, 28, 85.

Griffin, R.H. & Nunn, C.L. (2012) Community structure and the spread of infectious disease in primate social networks. Evolutionary Ecology, 26, 779–800.

Ilany, A. & Akçay, E. (2016) Social inheritance can explain the structure of animal social networks. Nature Communications, 7, 12084.

Ives, A., Dennis, B., Cottingham, K. & Carpenter, S. (2003) Estimating community stability and ecological interactions from time‐series data. Ecological Monographs, 73, 301–330.

Jarrett, J.D., Bonnell, T.R., Young, C., Barrett, L. & Henzi, S.P. (2018) Network integration and limits to social inheritance in vervet monkeys. Proceedings of the Royal Society B: Biological Sciences, 285, 20172668.

Josephs, N., Bonnell, T., Dostie, M., Barrett, L. & Henzi, S.P. (2016) Working the crowd: sociable vervets benefit by reducing exposure to risk. Behavioral Ecology, arw003.

Lubbe, A., Hetem, R.S., McFarland, R., Barrett, L., Henzi, P.S., Mitchell, D., Meyer, L.C., Maloney, S.K. & Fuller, A. (2014) Thermoregulatory plasticity in free-ranging vervet monkeys, Chlorocebus pygerythrus. Journal of Comparative Physiology B, 184, 799–809.

McFarland, R., Barrett, L., Boner, R., Freeman, N.J. & Henzi, S.P. (2014) Behavioral flexibility of vervet monkeys in response to climatic and social variability. American Journal of Physical Anthropology, 154, 357–364.

McFarland, R., Fuller, A., Hetem, R.S., Mitchell, D., Maloney, S.K., Henzi, S.P. & Barrett, L. (2015) Social integration confers thermal benefits in a gregarious primate. Journal of animal ecology, 84, 871–878.

Newman, M. (2010) Networks: an introduction. Oxford University Press, Oxford, UK.

